# Neutrophil: Airway Epithelial Interactions Result in Increased Epithelial Damage and Viral Clearance during RSV Infection

**DOI:** 10.1101/2019.12.20.885624

**Authors:** Yu Deng, Jenny A. Herbert, Elisabeth Robinson, Luo Ren, Rosalind L. Smyth, Claire M. Smith

**Affiliations:** Infection, Immunity and Inflammation, UCL Great Ormond Street Institute of Child Health, London, UK; Department of Respiratory Medical Centre, Chongqing Key Laboratory of Child Infection and Immunity, Children’s Hospital of Chongqing Medical University, China International Science and Technology Cooperation base of Child development and Critical Disorders, Ministry of Education Key Laboratory of Child Development and Disorders, Chongqing, 400014, China

**Keywords:** RSV, CILIA, NEUTROPHIL, INNATE IMMUNE RESPONSE, AIRWAY, EPITHELIUM, APOPTOSIS

## Abstract

Respiratory syncytial virus (RSV) is a major cause of paediatric respiratory disease. Large numbers of neutrophils are recruited into the airways of children with severe RSV disease. It is not clear whether or how neutrophils enhance recovery from disease or contribute to its pathology.

Using an *in vitro* model of the differentiated airway epithelium, we found that addition of physiological concentrations of neutrophils to RSV infected nasal cultures was associated with greater epithelial damage with lower ciliary activity, cilia loss, less tight junction expression (ZO-1) and more detachment of epithelial cells than seen with RSV infection alone. This was also associated with a decrease in infectious virus and fewer RSV positive cells in cultures after neutrophil exposure compared to pre-exposure. Epithelial damage in response to RSV infection was associated with neutrophil activation (within 1h), and neutrophil degranulation with significantly greater cellular expression of CD11b, MPO and higher neutrophil elastase and myeloperoxidase activity in apical surface medias compared to that from mock-infected AECs. We also recovered more apoptotic neutrophils from RSV infected cultures (>40%), compared to <5% in mock infected cultures after 4h.

The results of this study could provide important insights into the role of neutrophils in host response in the airway.

## INTRODUCTION

Respiratory syncytial virus (RSV) is the major viral cause of pulmonary disease in young infants and the elderly and is responsible for annual epidemics that cause considerable morbidity and mortality worldwide (1, 2). A growing body of evidence suggests that the virus initiates infection by targeting human ciliated epithelium lining the nasopharynx (3-6). Recent advances in cell culture have allowed us to explore the early effects of RSV infection on ciliated human respiratory epithelium and helped elucidate the mechanisms by which RSV causes disease.

Severe RSV disease is characterized by profound neutrophilic inflammation in the lungs: up to 85% of broncho-alveolar lavage (BAL) cells, from babies with bronchiolitis, are neutrophils (7). These cells are thought to play an important role in host defense during respiratory viral infections, but they have also been implicated in lung tissue damage in a variety of conditions including ARDS, acute lung injury, cystic fibrosis (CF) and COPD (8-10). In these and other conditions, it may be that neutrophils recruited to the airways as part of host defense contribute to tissue damage and exacerbate disease. Currently little is known about the role of neutrophils in the airways during RSV disease. Studies using animal models (11) (e.g. human RSV infection in mice, sheep, and cotton rats) lack sensitivity and do not fully recapitulate the disease in man. For example, only 18-27% of BAL cells recovered from human RSV infected calves are neutrophils, 3-14% in mice (12, 13) and <5% in cotton rats (14).

In the human airway, infiltrating neutrophils directly interact with inflamed epithelial surfaces and have adapted mechanisms to respond to RSV infected airway epithelial cells. However, few studies that have investigated neutrophil function during this interaction. The primary aim of this study was to evaluate the possible injury caused by neutrophils to RSV infected ciliated cell cultures, to evaluate the early changes in neutrophil function during RSV respiratory infection and to assess whether neutrophils had an anti-viral effect.

## METHODS

### Subjects and ethical considerations

Human nasal epithelial cells and/or peripheral blood samples were obtained from different (heterologous) adult healthy control donors who had no history of nasal or respiratory disease. Epithelial cells were collected at least 3 months prior to neutrophil collection. None of the subjects were taking medications or had reported a symptomatic respiratory infection in the preceding 6 weeks. All samples were obtained with the individual’s permission and with ethical approval by the UCL Research Ethics Committee (ref 4235/002).

### Viral infection of primary epithelial cell cultures

Primary human nasal airway epithelial cells (nAECs) were grown to a ciliated phenotype on 0.4µm pore Transwell inserts (Corning) at air–liquid interface (ALI) as described previously (3). Recombinant GFP tagged RSV A2 strain was kindly provided by Fix et al, 2011 (15) and propagated using HEp-2 cells (MOI 0.1) for 3-5 days in Opti-MEM. Virus was purified as described previously (16), collected in BEBM (Life Technologies) and frozen at −80°C. Aliquots of RSV were thawed immediately prior to use. For ALI culture infection, the apical surface of the ALI cultures was rinsed with medium (BEBM) and 200µL viral inoculum (MOI: 1) in BEBM was applied to the apical surface for 1h at 37°C and then removed (see Supplementary **Figure S1**). Mock wells received BEBM alone. All cells were fed basolaterally with fresh ALI medium prior to infection. The infection continued for 24h and 72h.

### Neutrophil isolation and purification

Neutrophils were isolated from peripheral venous blood using a Percol density gradient as described previously (17) and purified using the EasySep™ Human Neutrophil Enrichment Kit (STEMCELL Technologies) as per the manufacturer’s instructions (18). Neutrophils were resuspended in Hanks balanced salt solution (HBSS-) (Life Technologies) and FACS analysis was performed using Anti-Human CD49d-APC (BioLegend, 304307) and Anti-Human CD66a/c/e AlexaFluor-488 (BioLegend, 342306) antibodies to confirm purity. Neutrophils were counted using a haemocytometer.

### Neutrophil airway epithelial cell co-culture

Neutrophils (5×10^5^ in 100μL HBSS+) were added to the top (apical) chamber of Transwell inserts containing either RSV or mock-infected ciliated nAECs and incubated at 37°C+5% CO2. After 1h or 4h incubation, neutrophils and apical surface media from the top chamber of Transwell were collected and each membrane was washed once with 0.1mL HBSS+. The apical surface media and washing medium were pooled, spun and stored at −80°C (see Supplementary file). The cell pellet was resuspended in 500µl FACS buffer (PBS (Ca2+Mg2+ free), 0.5% bovine serum albumin (BSA), 2.5mM EDTA) for further analysis.

### Determination of neutrophil degranulation

Degranulation was assessed by measuring neutrophil elastase (NE), myeloperoxidase (MPO and matrix metalloproteinase-9 (MMP-9) in the apical surface media. The amount of NE or MPO in apical surface media was measured using commercial activity assay kits (Cayman, USA). MMP-9 release was measured using commercial ELISA kit (Biolegend, USA). All protocols were according to the manufacturers’ instructions.

### Neutrophil CD11B and MPO expression

Neutrophil activation was determined by measuring the cell surface protein expression levels of CD11B and MPO. All neutrophils were centrifuged at 1400rpm, 5mins and washed once in 500µl FACS buffer (PBS (Ca2+Mg2+ free), 0.5% BSA, 2.5mM EDTA). The cell pellet was then resuspended in 50µl (1/50 dilution in FACS buffer) of TruStain FcX blocker antibody (BioLegend) and incubated at 4°C for 10minutes, then washed in FACS buffer as above. Cells were resuspended in 50µl FACS buffer plus 1/250 dilution PE anti-Human CD11B conjugate (50-0118-T100, Insight Biotechnology) and 1/50 dilution Anti-MPO-APC human antibody (130-107-177, clone:REA491, Miltenyl Biotec) and incubated at 4°C for 20 minutes in the dark. Unstained and single antibody controls confirmed no cross reactivity. Cells were washed once in FACS buffer, resuspended in 1% (w/v) PFA and stored at 4°C. Directly prior to running, samples were centrifuged (1400rpm) and resuspended in FACS buffer.

Samples were analysed using a Beckton Dickenson LSR II flow cytometer and FlowJo v10.0 FACS analysis software. Neutrophils were first identified as being CD11B positive (PE+). Using this population the mean fluorescence intensity for PE and APC was calculated.

### AnnexinV/PI apoptosis assay

AnnexinV apoptosis detection kit (Miltenyl Biotec) was used to carry out this assay. Neutrophils collected from the cell pellet (described above) were resuspended in 50µl FACS buffer plus 1:250 dilution APC anti-Human CD11B conjugate (Insight Biotechnology) and incubated at 4°C for 20 minutes in the dark. Cells were washed once in FACS buffer, resuspended in 50µl Annexin binding buffer with 0.3µl of Annexin V for 15 minutes in the dark at room temperature. They were then flash stained with propidium iodide (PI), 10µl in 900µl of Annexin binding buffer and analysed on a FACSCalibur flow cytometer. Unstained and Annexin V or PI single stained controls confirmed no cross reactivity. At least 10,000 events were collected. Neutrophils were first identified as being CD11B positive (PE+).

### CBF and beat pattern

Beating cilia were observed via an inverted microscope system (Nikon TiE; Nikon, UK) equipped with an incubation chamber (37°C, 5% CO2) as previously described (3). To determine ciliary beat frequency (CBF), videos were recorded using a ×20 objective using a CMOS digital video camera (Hamamatsu) at a rate of 198 frames per second, and image size of 1024×1024 pixels. CBF (Hz) was calculated using ciliaFA software (19). The number of motile ciliated cells in each sample area was counted (motility index). The dyskinesia index was calculated as the percentage of dyskinetic ciliated cells (those that displayed uncoordinated motile cilia or those that beat with a stiff, flickering or twitching motion) relative to the total number of motile ciliated cells.

### Immunofluorescence microscopy

Following fixation, cells were washed three times with PBS, treated with PBS containing 0.1% Triton X-100 for 10 min to permeabilize the cells. Cells were incubated with 5% FCS in PBS for 0.5h at room temperature to block nonspecific interactions, and washed again three times with PBS. All subsequent antibody incubations were carried out in 5% FCS in PBS + 0.1% Triton X-100. Reagents used in this study were rabbit anti-ZO-1 polyclonal antibody (1:200, sc-5562, Santa Cruz) and mouse anti-acetylated α-tubulin monoclonal antibody (6-11B-1; 1μg/mL; Sigma). Primary antibody incubations were carried out in a humidified chamber overnight at 4°C, followed by three washes with PBS. Detection of primary antibodies was carried out for 1h using the following reagents: fluorescein FITC (Sigma #F2012) conjugated rabbit anti-mouse (1:64) or AlexaFluor 594-conjugated rabbit anti-donkey antibody (1:250; Invitrogen, Paisley, UK). All secondary antibodies had been tested and found to be negative for cross-reactivity against human epithelial cells. Following three washes in PBS, DNA was stained with Hoechst 33258. After a final wash in distilled water, the insert was cut from the support and mounted under coverslips in 80% (v/v) glycerol, 3% (w/v) n-propylgallate (in PBS) mounting medium. Images were captured with confocal laser microscope (Zeiss Observer Z.1) using a 40x water immersion objective. The pinhole was set at 1 airy unit (AU). For Z-stack images the slice thickness was 1μm.

### Statistical analysis

Statistical analysis was performed using GraphPad Prism 5 (GraphPad, San Diego, CA, USA). Differences between mock and RSV infected group were analysed using paired t-tests or for multiple group comparisons a paired two-way ANOVA for multiple comparisons with a Bonferroni correction was used (GraphPad Prism v5.0).

## RESULTS

### Neutrophils enhance ciliated epithelial layer disruption and ciliary loss

We found that neutrophils interacted with motile ciliated nAECs almost immediately after introduction and gathered together to in clusters (**Figure 1A**). After 1h incubation with ciliated nAECs infected with RSV for 24h and 72h, neutrophils were shown to decrease epithelial expression of ZO-1 (a marker of epithelial cell tight junction proteins) when compared to mock-infected cultures (P=0.002). This reduction in ZO-1 expression was also significantly lower than RSV infected cells without neutrophils (**Figure 1B and C**) at 72h (P=0.001), but not 24h post infection (P=0.07). We also found that addition of neutrophils was associated with greater nAEC loss after 1h, with significantly fewer RSV-infected nAEC nuclei remaining attached to the membrane insert at 24h post-RSV infection compared to the respective mock-infected control (P=0.006) and compared to RSV-infected nAEC without neutrophils (P<0.0001) (**Figure 1D**). ZO-1 staining and numbers of epithelial cells were similar whether neutrophils were added to cultures infected for 24 or 72 hours.

**Figure 1.**
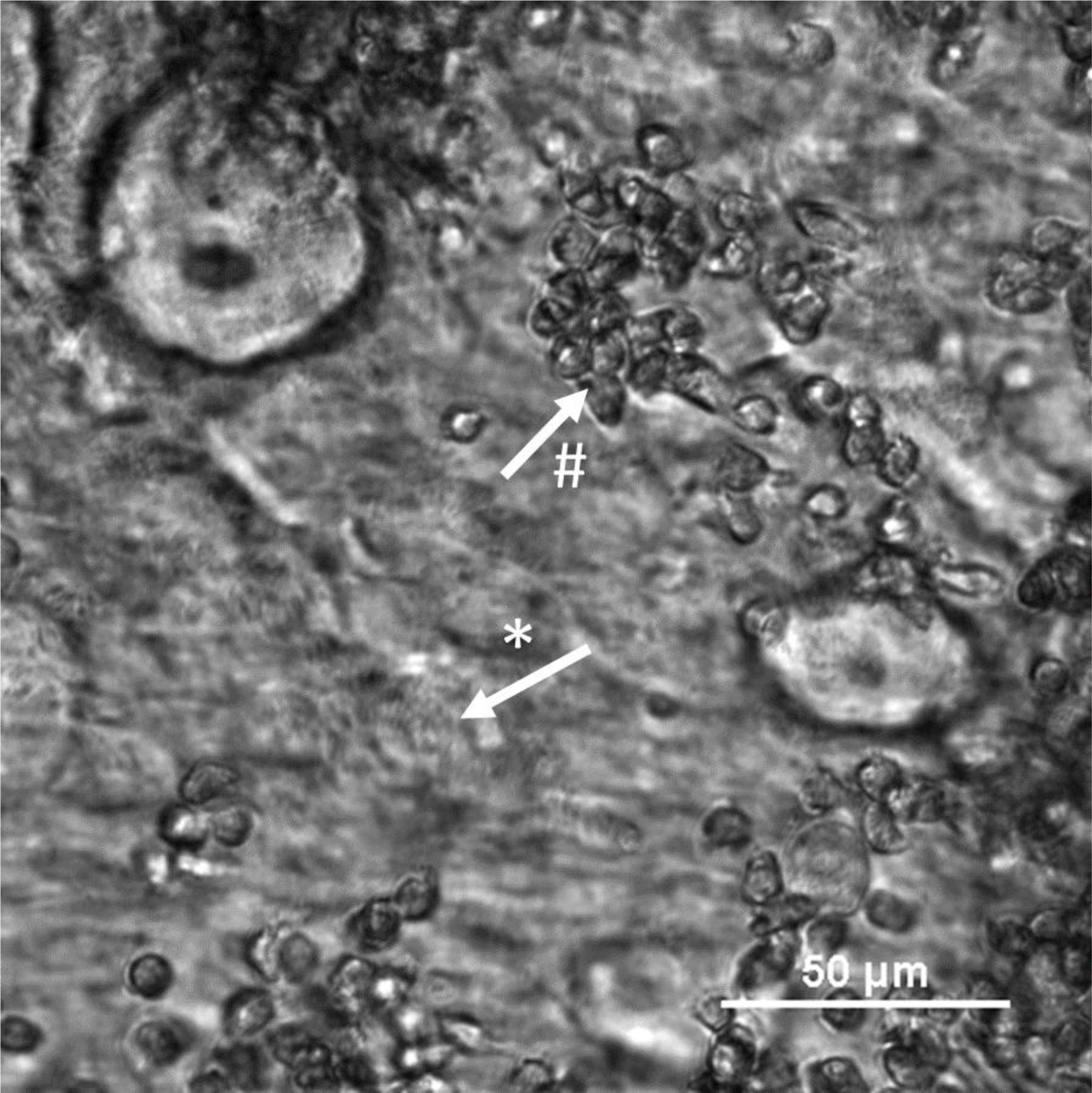
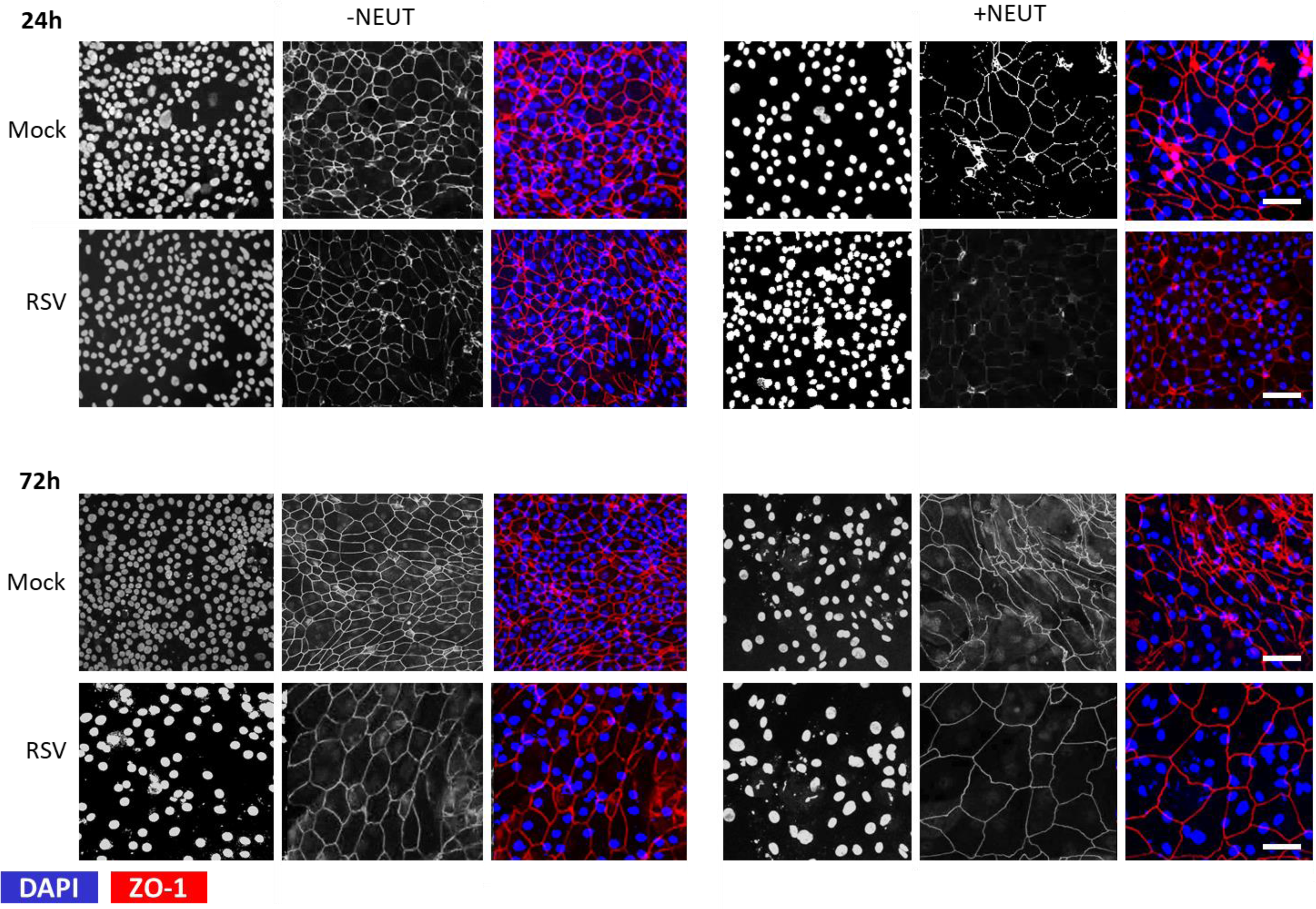
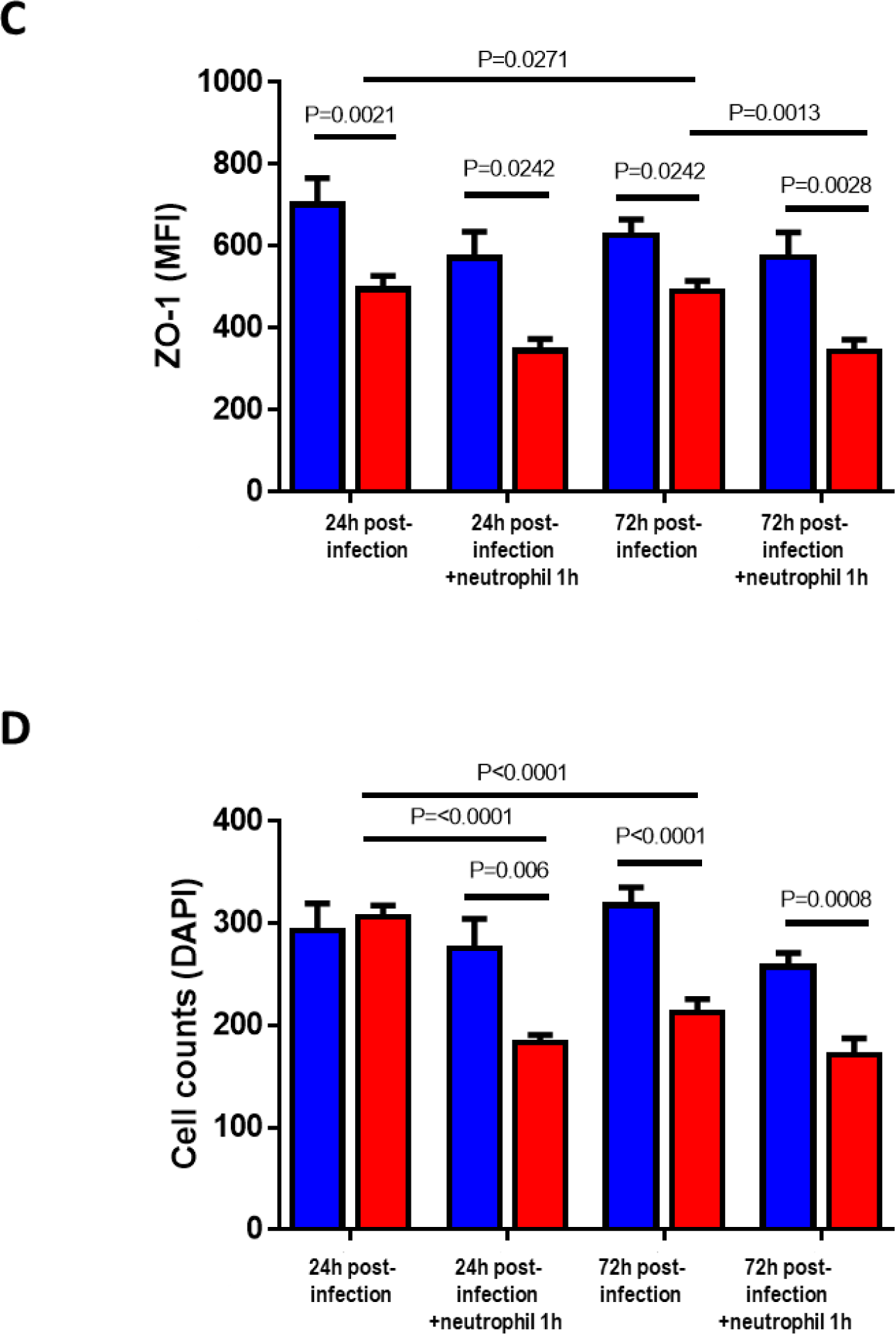
Epithelial damage caused by neutrophil exposure to RSV infected ciliated epithelial cells. The effect of respiratory syncytial virus (RSV) and neutrophil exposure on the number of α-tubulin positive cells. (**A**) Bright-field image of an RSV infected culture after 4h co-culture with neutrophils showing ciliated epithelial cells (*) and neutrophil clusters (#). Scale bar shown. (**B**) Representative confocal images of RSV-infected human nasal ciliated epithelial cells with and without neutrophil exposure for 1h. Cells were stained with antibodies against ZO-1 to detect tight junctions and DAPI to detect epithelia nuclei. (**C**) Mean fluorescence intensity of ZO-1 of ciliated airway epithelial cells infected with mock (blue bars) or RSV (red bars) for 24h or 72h cells and after 1h neutrophils (n=6 epithelial donors, 6 heterologous neutrophils). Bars represent the mean±SEM. P values show a significant differences. **(D)** The number of epithelial cells attached to membrane inserts after neutrophil exposure. Epithelial cells were quantified by counting the DAPI stained nuclei >50µm^2^ in area using ImageJ, the mean (SEM) number of epithelial cells from all images is shown (n=5 images per donor, 3 epithelial donors with heterologous neutrophils).

As RSV has been shown to target ciliated cells for infection, we were especially interested in how ciliary activity is altered during neutrophil interactions with RSV infected epithelial cells. We found that ciliary beat frequency was unaffected by RSV infection or exposure over the entire study period (**Table 1**). However, RSV infection led to a higher proportion of dyskinetic cilia (**Table 1**). The mean dyskinesia index was significantly higher at 24h post-RSV infection (25.71±1.14%) and after 72h of RSV infection (42.38±2.44%) compared with mock-infected controls (8.44±0.20%) and (10.50±0.35%), respectively (P<0.001).

**Table I.** The ciliary beat frequency, dyskinesia index and motility index of healthy nasal respiratory epithelial cells in pseudo-stratified air–liquid interface (ALI) cultures infected with RSV A2 for 24h or 72h and then co-cultured for 1 or 4h with human neutrophils.

|  | Hours post-neutrophil | Mean beat frequency (Hz) |  | Mean dyskinesia index (%)<sup>a</sup> |  | Mean motility index<sup>b</sup> |  |
| --- | --- | --- | --- | --- | --- | --- | --- |
|  |  | Control | RSV-infected | Control | RSV-infected | Control | RSV-infected |
| **24h** | 0 | 10.84±0.28 | 11.64±0.16 | 8.439±0.20 | **25.71±1.13<sup>\*</sup>** | 63.51±1.66 | 61.4±1.67 |
|  | 1 | 12.65±1.00 | 12.56±0.48 | 9.582±0.28 | **25.77±1.15<sup>\*</sup>** | 68.37±2.43 | 60.86±1.82 |
|  | 4 | 12.23±1.07 | 12.08±0.74 | 10.04±0.84 | **26.91±2.41<sup>\*</sup>** | 65.1±1.73 | 61.63±2.35 |
| **72h** | 0 | 12.76±0.35 | 13.65±0.28 | 10.50±0.35 | **42.38±2.44<sup>\*</sup>** | 70.78±1.97 | **57.6±1.89<sup>\*</sup>** |
|  | 1 | 12.12±1.59 | 11.29±0.98 | 11.03±0.45 | **49.45±2.03<sup>\*</sup>** | 66.94±2.93 | **50.41±2.77<sup>\*#</sup>** |
|  | 4 | 11.71±1.30 | 10.87±1.07 | 10.23±0.59<sup>a</sup> | **44.96±2.62<sup>\*</sup>** | 71.02±2.36 | 65.4±3.71 |

<sup>a</sup>Motility index, number of motile ciliated cells per sample area of ~4.2mm<sup>2</sup>. <sup>a</sup>Dyskinesia index, percentage of abnormally beating cilia amongst all cilia examined..
Data expressed as mean ± SEM. Values highlighted in boldface were significantly different = <sup>\*</sup>p<0.05 when compared with Mock. # p<0.05 when compared to 0h.

Addition of neutrophils to RSV infected nAECs did not further enhance ciliary dyskinesia at any time point or infection condition tested (**Table 1**). Representative slow motion videos showing ciliary beating are show in supplementary videos **VIDEO 2-5**. We found no significant difference in mean fluorescence intensity for α-tubulin staining (cilia tubulin marker) under fluorescence microscopy examination at 24h post-RSV infection, compared to mock-infected cells or following 24h RSV infection with and without neutrophil exposure (**Figure 2A/B**). However, we found that at 72h post-RSV infection, the mean fluorescence intensity for α-tubulin was almost half that of the mock-infected group with neutrophils (P=0.014) (**Figure 2B**). This loss α-tubulin staining correlated with a loss in the number of motile cilia, observed by light microscopy (referred to as the mean motility index) at 72h post-RSV infection (57.6±1.89%), which was less compared with the mock-infected controls (70.8±1.97%) (P<0.05). Exposure to neutrophils for 4h lowered the mean motility index of ciliated cells infected with RSV for 72h (50.41±2.77%) compared to pre-neutrophil time point (P<0.05) (**Table 1**). As is shown in **Figure 2C**, the distribution of ciliary beat frequency of the same field of view before and after addition of neutrophils produced a similar mean CBF of around 10.2-10.9Hz, but fewer areas (ROI) from the RSV infected ciliated nAECs (bottom right panel of **Figure 2C**) show active beating cilia (defined as >3Hz) when co-cultured with neutrophils.

**Figure 2.**
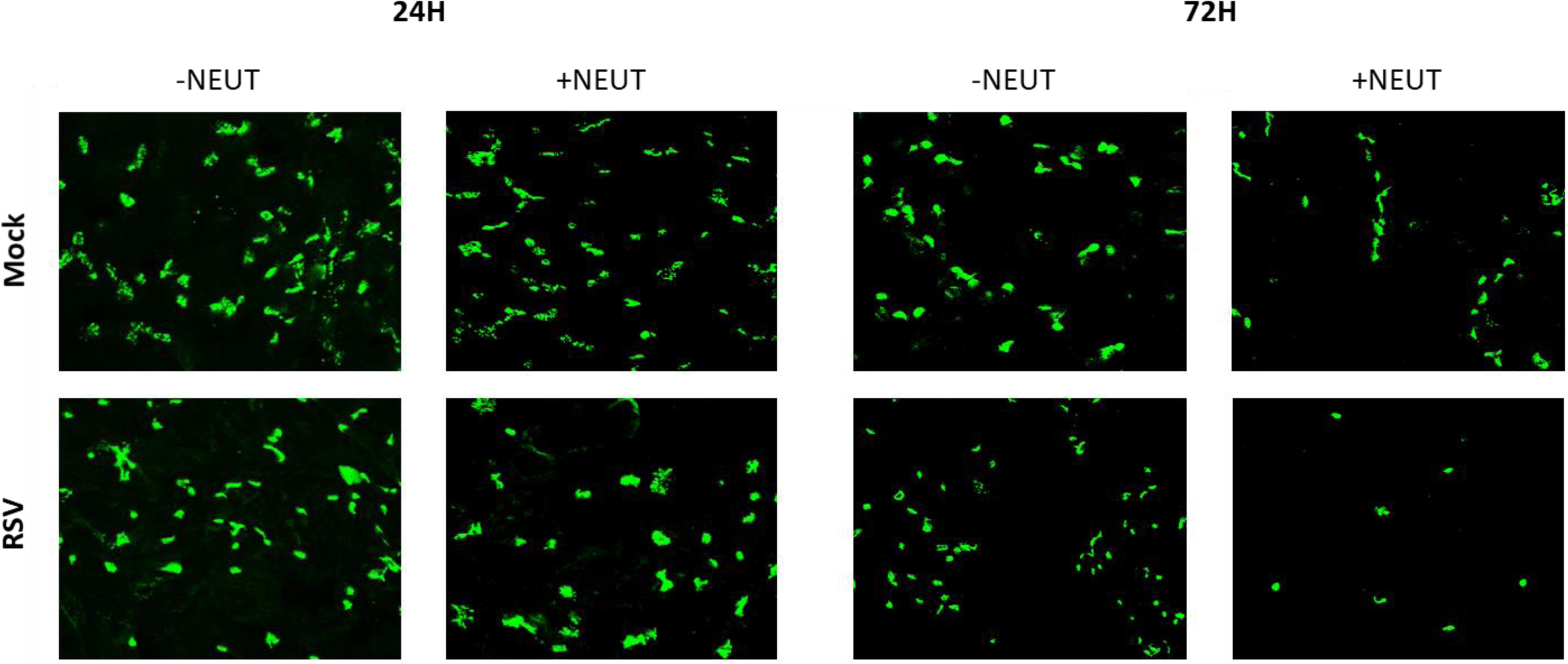
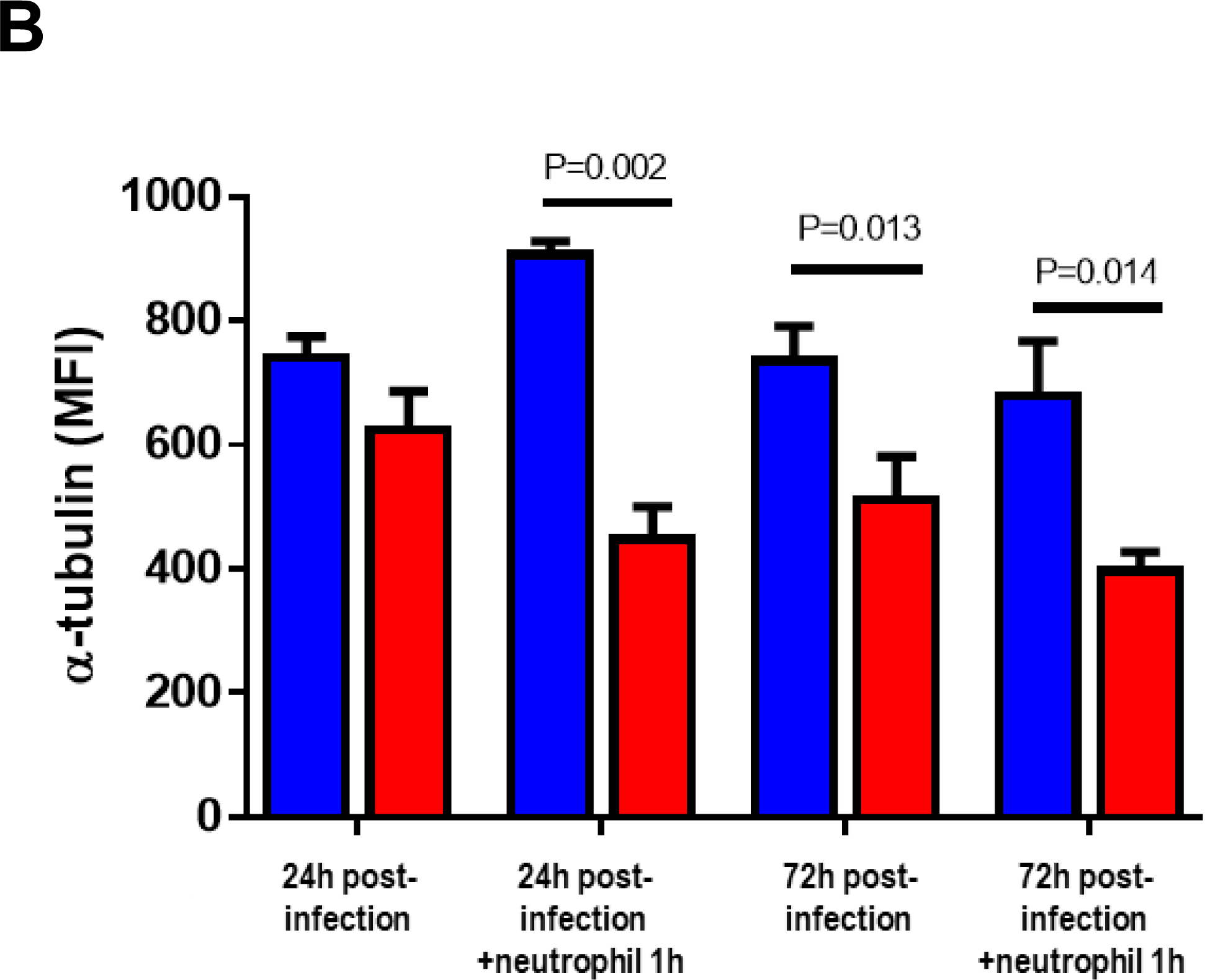
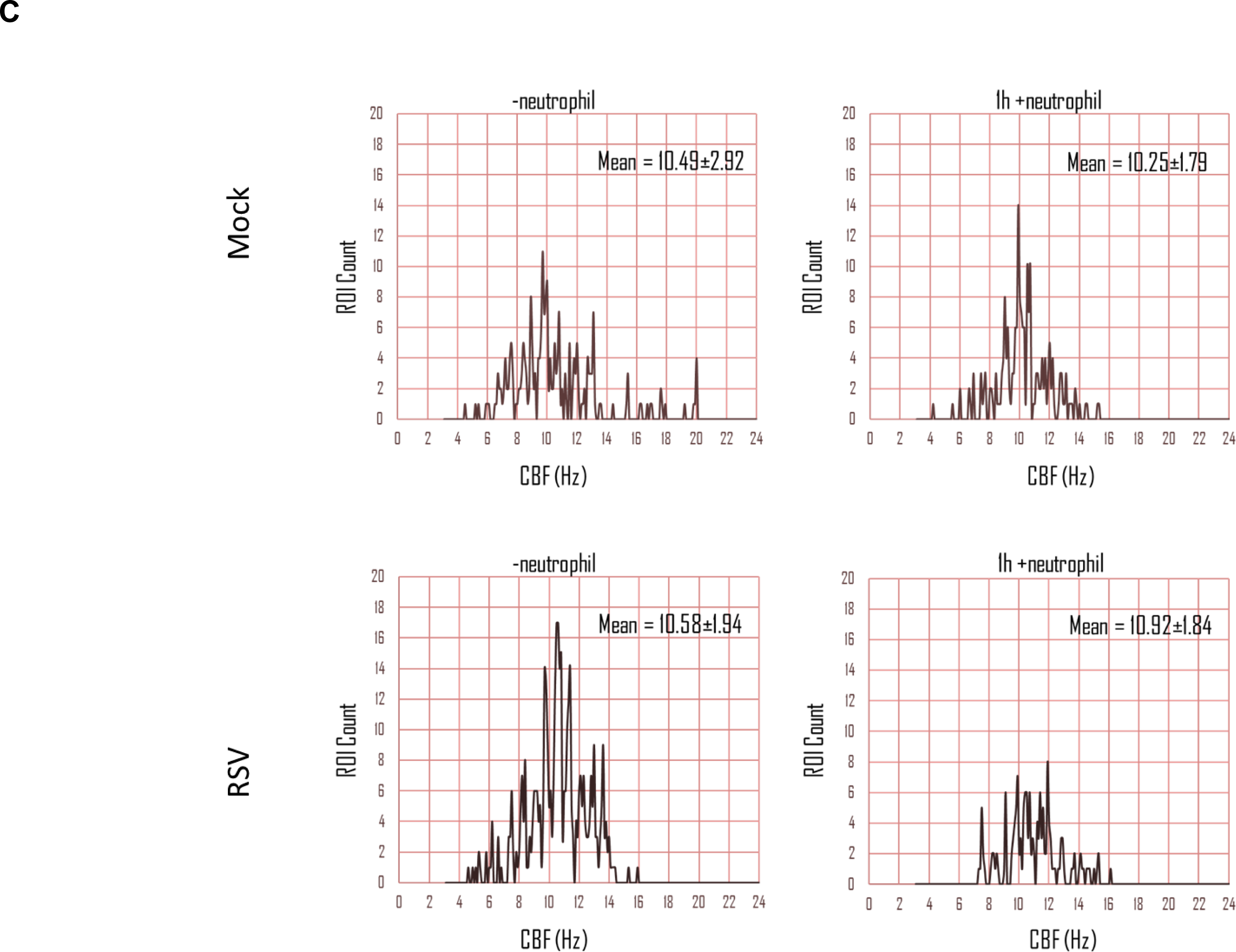
The number of ciliated epithelial cells decreases after neutrophil exposure. (**A**) (**B**) Representative confocal images of RSV-infected human nasal ciliated epithelial cells following neutrophil exposure for 1h. Cells were stained with antibodies against acetylated tubulin to detect the ciliary axonemal microtubules (green). (**C)** The number of ciliated cells present on membrane inserts were quantified after 24h or 72h infection and 1h neutrophil exposure. The level of α-tubulin staining was used as a measure of cilia present (n=5 images per donor, 3 epithelial donors with heterologous neutrophils). Bars represent the mean±SEM for mock-infected (blue bars) or RSV infected (red bars) cultures. (**G**) Histograms of the frequency distribution of ciliary beat frequency from 1600 regions of interest taken from a representative field of view.

### Incubation with neutrophils reduces the amount of infectious virus in the culture

After 4h after neutrophil exposure, we detected a significantly lower infectious virus with a RSV titre of 2.4×10^4^pfu/ml recovered from cultures infected with RSV for 72h without neutrophils, compared to 2.8×10^3^pfu/ml in cultures infected with RSV for 72h with neutrophils (P<0.05) (**Figure 3A**). We assessed the numbers of RSV infected nAECs by measuring the mean fluorescence intensity of GFP, a marker of active cytoplasmic RSV replication. Using fluorescence microscopy (**Figure 3B**), we found that GFP expression was significantly lower 1h after exposure to neutrophils compared with those cells without neutrophils at both 24h and 72h post infection (P<0.05) (**Figure 3B and C**).

**Figure 3.**
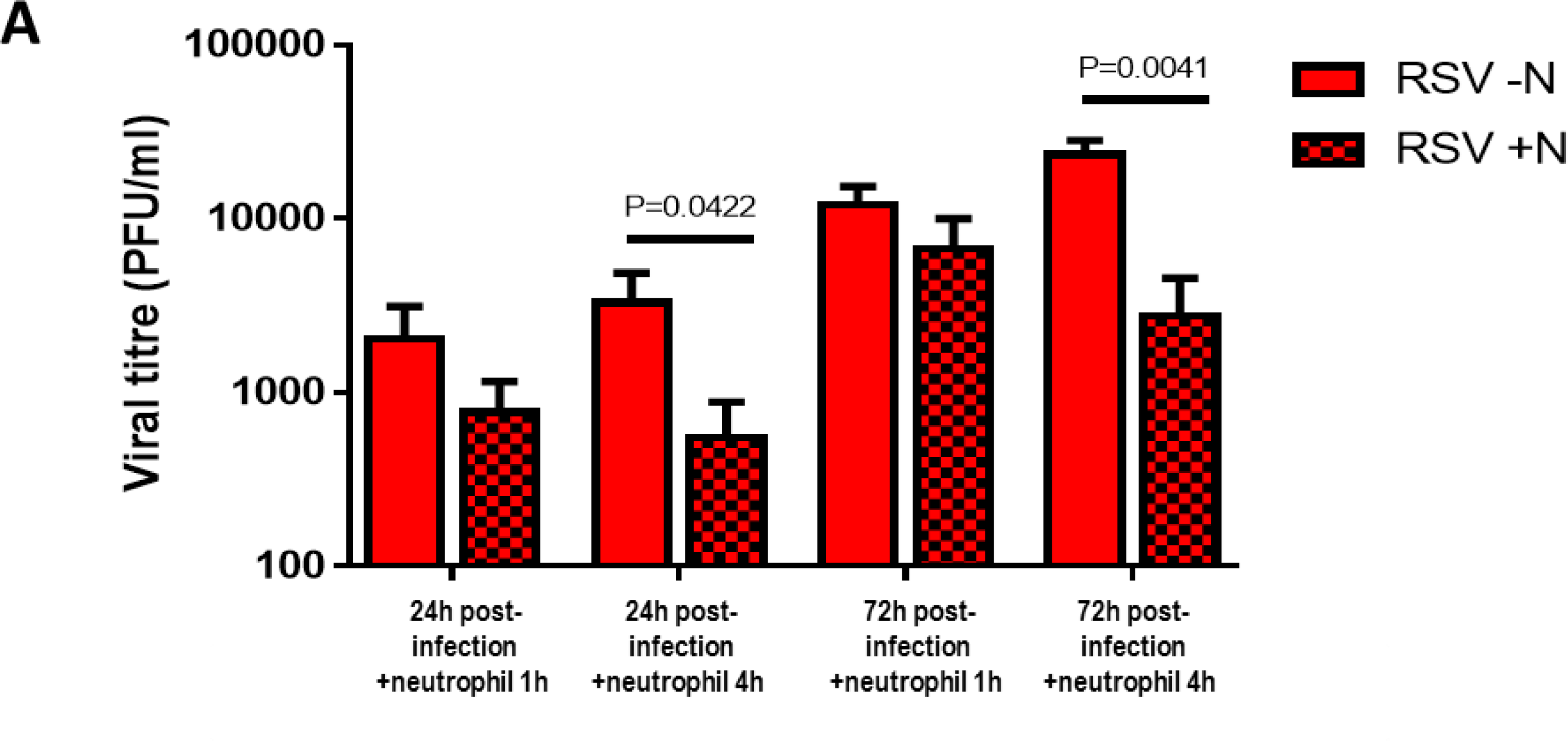
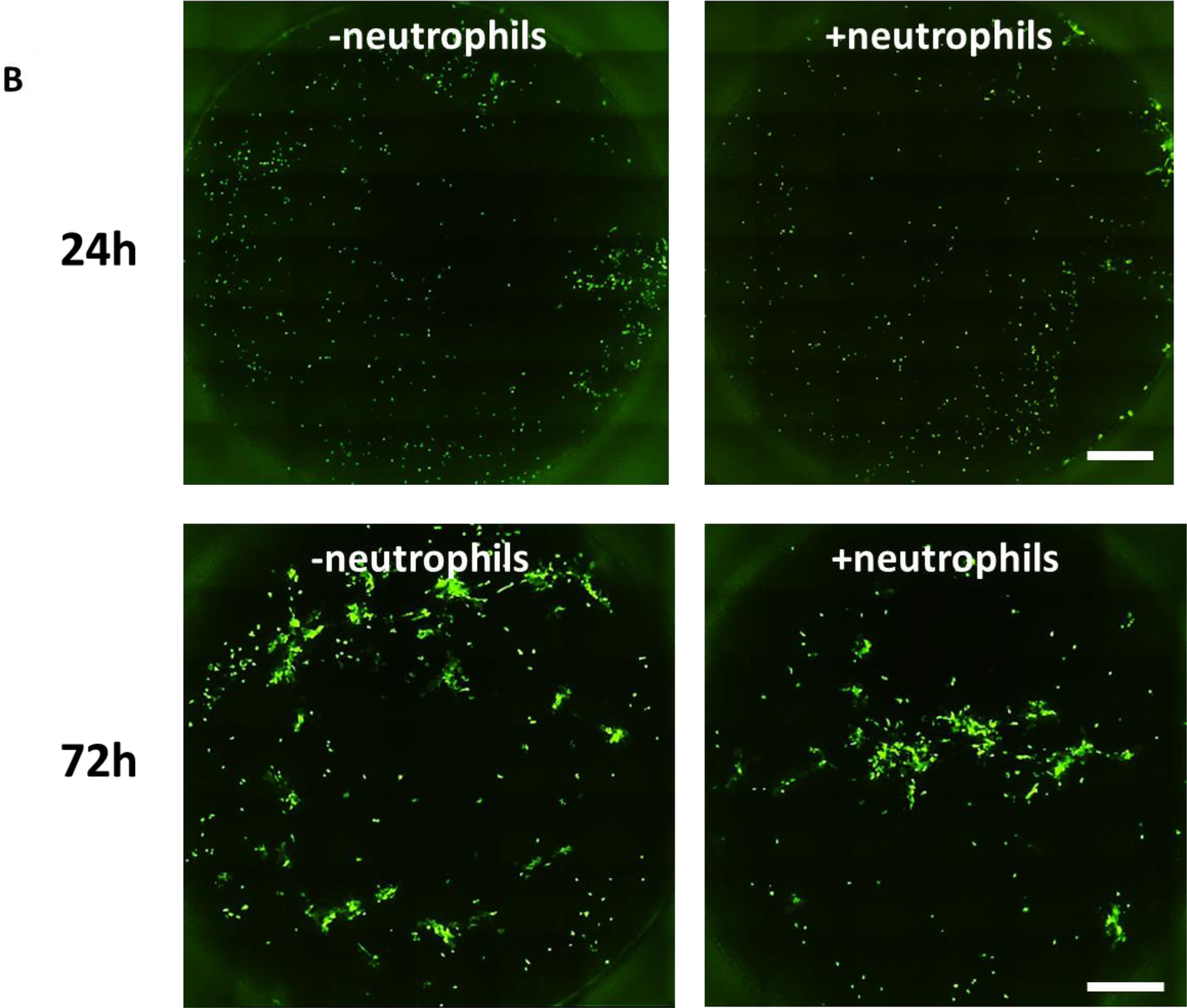
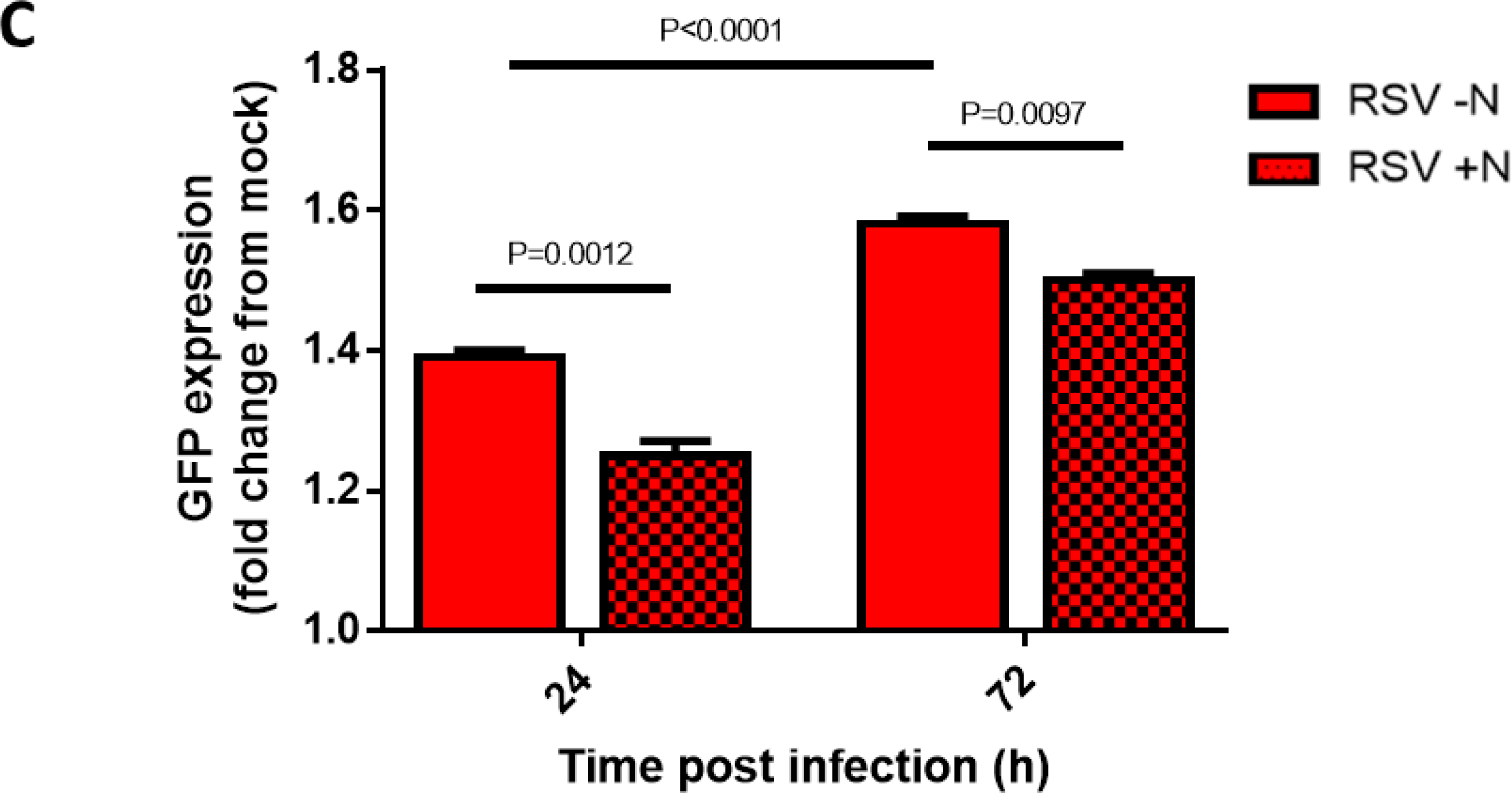
The amount of infectious RSV is decreased in infected ciliated epithelial cells exposed to neutrophils. (**A**) The amount of infectious virus present in whole wells following neutrophil co-culture as measured by plaque forming units (pfu/ml). (**B**) Whole well scan of a representative membrane insert infected with RSV for 24h or 72h cells before (left panel) and 1h after (right panel) neutrophil exposure. Each white spot indicates an RSV infected epithelial cell. (**C**) Mean fluorescence intensity of GFP-RSV infected AECs for 24h or 72h cells after 1h without (clear bars) and with (spotted bars) neutrophils. Bars represent the mean ± SEM (n=6 epithelial donors with heterologous neutrophils). P values show a significant differences between the matched pre and post neutrophil exposure time points.

### Incubation with RSV infected ciliated epithelial cells increases neutrophil apoptosis

Using flow cytometry (**Figure 4A**) we showed that, at 24h post RSV-infection, there was no difference in the percentage apoptotic (PI^lo^ AnnexinV^hi^) neutrophils recovered from RSV infected nAECs (11.1±5.1%) compared to 4.8±2.7% in the mock-infected co-cultures (**Figure 4B)**. However, at 72h post-infection, we detected significantly more apoptotic neutrophils after exposure to RSV infected nAECs with 46.7±11.4% compared to 6.2±0.9% in the mock-infected co-cultures (P<0.0001) (**Figure 4B)**. Apoptosis appeared to be the dominant form of cell death as we did not detect an increase in dead (PI^hi^ AnnexinV^lo^) neutrophils at any time point or test condition. After 4h exposure to epithelial cells infected with RSV for 72h we detected 6.43±7.2% dead neutrophils compared to 5.5±7.3% in the mock control (P>0.99) (**Figure 4B**).

**Figure 4.**
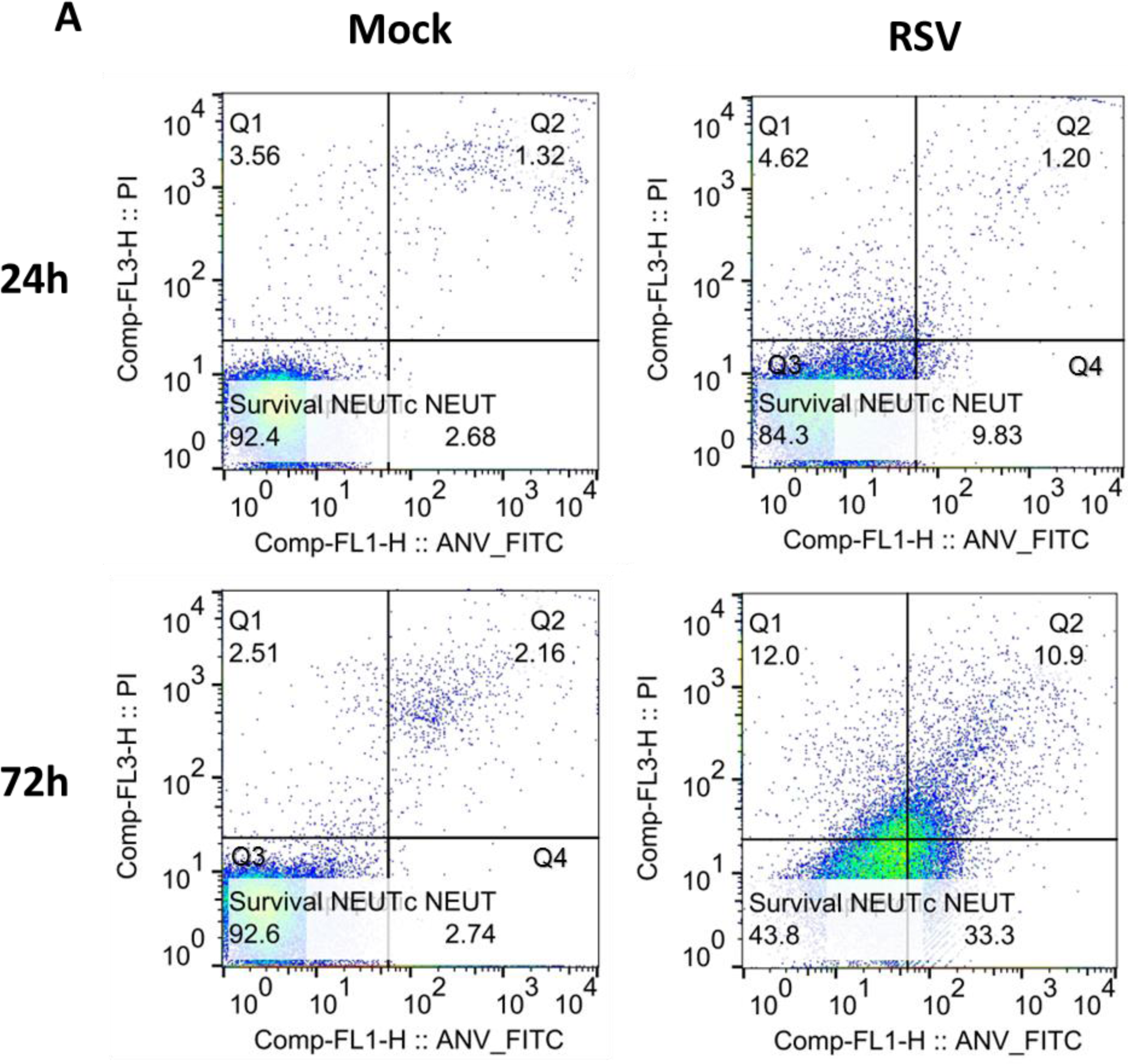
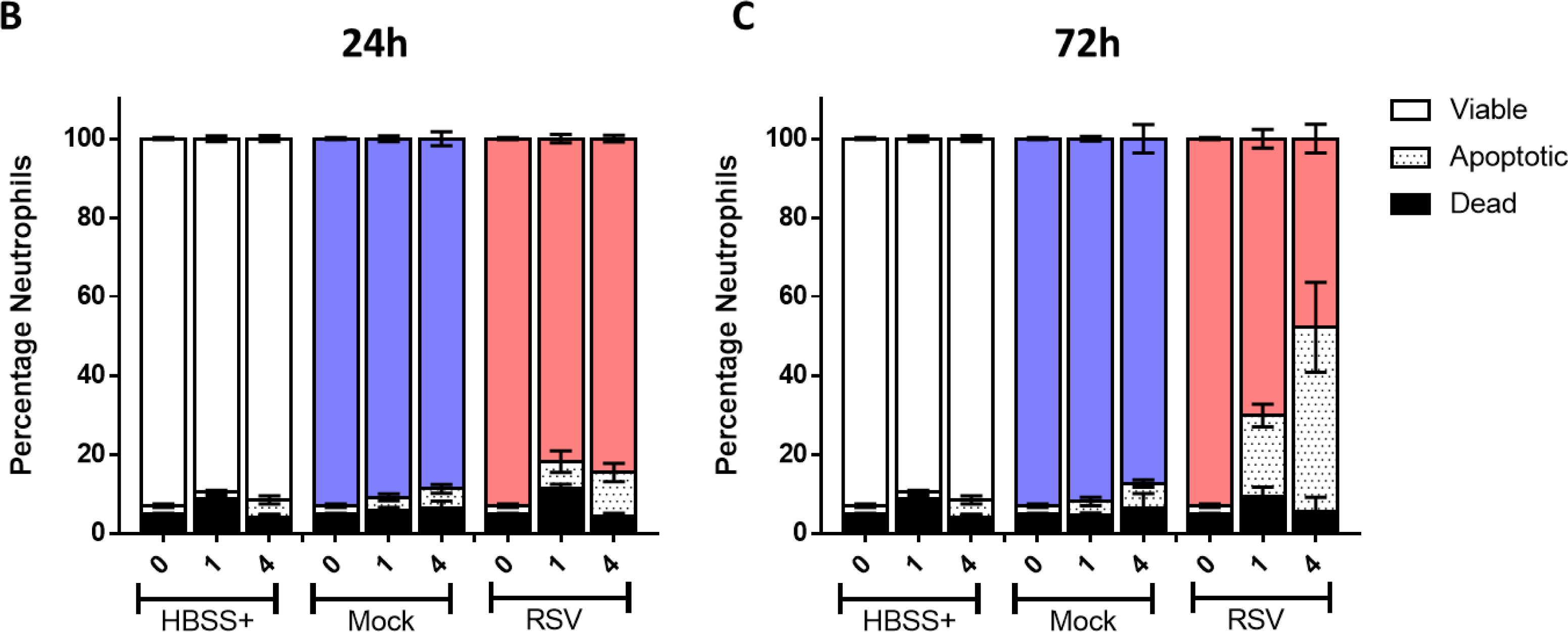
Neutrophil viability and apoptosis following exposure to RSV infected ciliated epithelial cells. **(A)** Viability was calculated by exclusion of propridium iodide (PI^lo^) and apoptosis was determined by expression of AnnexinV (FITC^i^) and determined by flow cytometry. Neutrophils were first identified using a CD11b (PE) positive gate. Using this proportion of the population with fluorescence intensity of PI and/or FITC fluorescence was calculated. Q1 = PI^hi^ANV^lo^, Q2 = PI^hi^ANV^hi^ (Q1+Q2=dead neutrophil), Q3 = PI^lo^ANV^lo^ (viable neutrophils), Q4 = PI^lo^ANV^hi^ (apoptotic neutrophils). (**B**) Bars show mean±SEM of n=4-7 epithelial donors with heterologous neutrophils for HBSS (white), mock-infected (blue bars) or RSV infected (red bars) cultures. Statistical significance is shown.

### Incubation with RSV infected ciliated epithelial cells increases neutrophil expression of CD11B and MPO and augments degranulation

Incubation with ciliated epithelial cells RSV infected for 24h led to significantly greater expression of CD11B on neutrophils after 4h, but not 1h, compared to neutrophils that were incubated with mock infected ciliated epithelial cells with a mean (±SEM) fluorescence intensity of 1.3×10^3^±1.21×10^2^ compared to 9.7×10^2^±3.9×10^1^ respectively after 4h (P=0.048) (**Figure 5A**). At 72h post infection, incubation with ciliated epithelial cells RSV infected led to significantly greater expression of CD11B on neutrophils after 1h and 4h, compared to neutrophils that were incubated with mock infected ciliated epithelial cells with a MFI of 1.7×10^3^±3.6×10^2^ compared to 8.9×10^2^±8.2×10^1^ respectively after 4h (P=0.0002) (**Figure 5A**). Incubation with epithelial cells infected for 24h showed significantly greater expression of MPO on neutrophils after 1h and 4h of incubation, compared to neutrophils that were incubated with mock infected ciliated epithelial cells with a mean (±SEM) fluorescence intensity of 6.5×10^2^±1.1×10^2^ compared to 2.5×10^2^±1.2×10^1^ respectively after 24h infection (P<0.0001) (**Figure 5B)** with similar findings at 1h (P=0.722) and 4h (P<0.0001) after 72h RSV infection.

**Figure 5.**
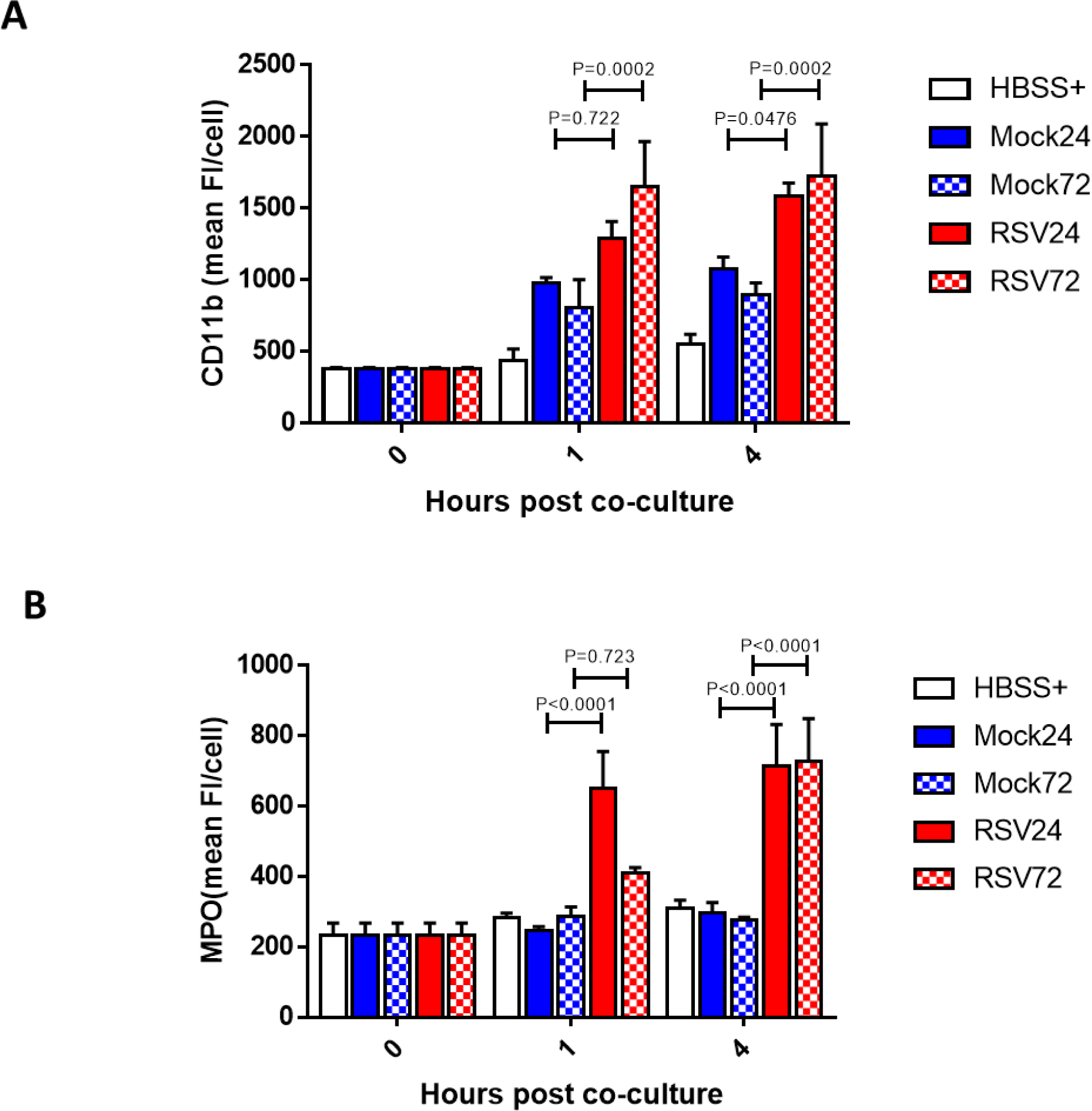
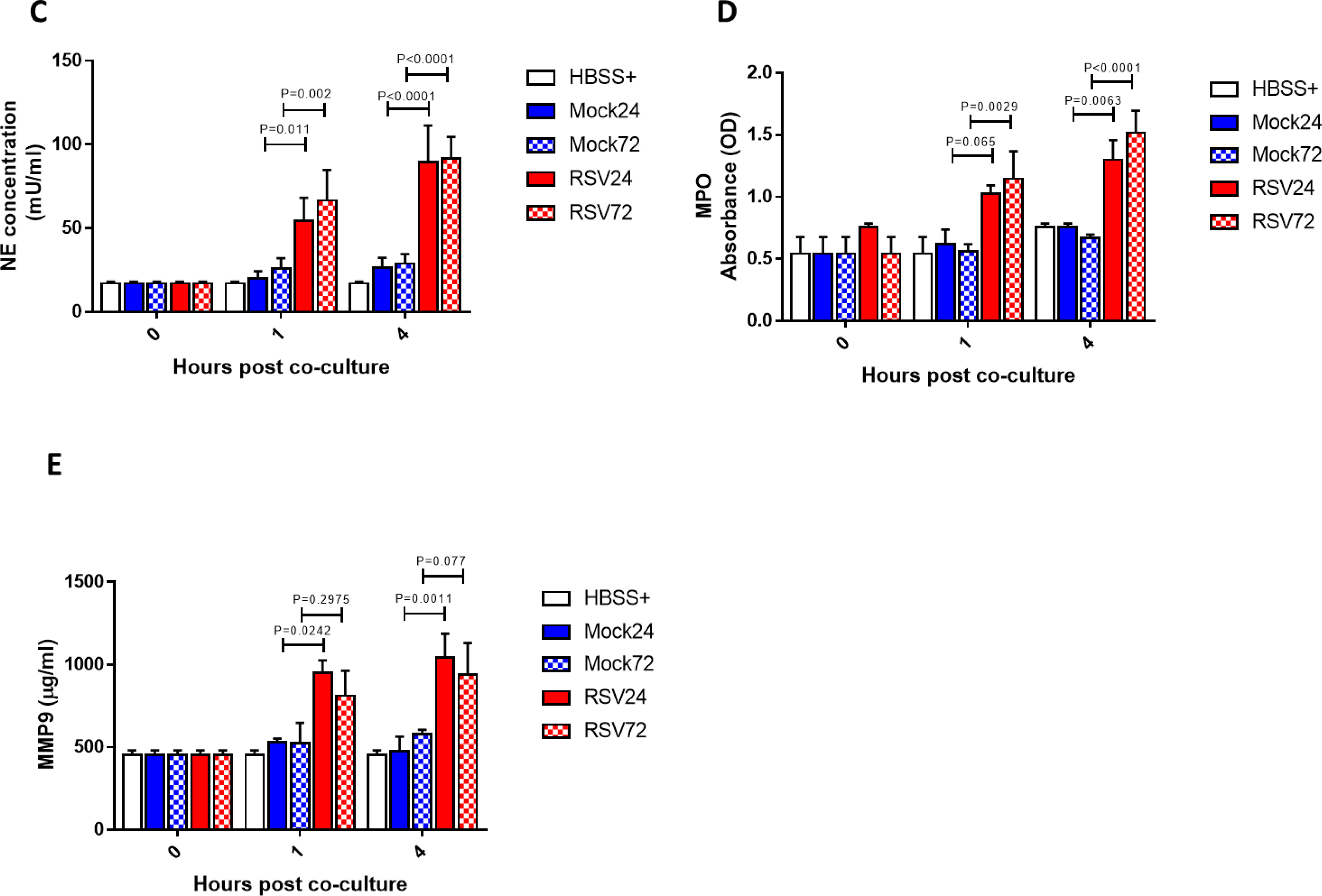
Neutrophil activation and release of neutrophil derived products and neutrophil activation after exposure to RSV infected ciliated epithelial cells. (**A**) Neutrophils in exposure to RSV infected epithelial cells infected for 24 (plain bars) or 72 hours (checkered bars) show increased cell surface expression of CD11b compared to those exposed to mock infected epithelial cells. (**B**) Neutrophils exposed to RSV infected epithelial cells infected for 24 (plain bars) or 72 hours (checkered bars) show increased cell surface expression of MPO compared to those exposed to mock infected epithelial cells. Activation marker expression was calculated by staining for cell surface expression of CD11b (PE^hi^) or MPO (APC^+^) and determined by flow cytometry. Neutrophils were identified using a PE positive gate. Using this population the geometric mean fluorescence intensity of PE or APC fluorescence was calculated. Bars show mean ± SEM of n=5-8 epithelial donors with heterologous neutrophils. Statistical significance is shown. Apical surface media concentrations of (**C**) neutrophil elastase (NE) and (**D**) myeloperoxidase (MPO) and (**E**) MMP9 were measured after exposure to mock or RSV infected (24 (plain bars) or 72 hours (checkered bars)) ciliated AECs at 1 and 4 hours. For all, bars represent the mean±SEM of n=5 epithelial donors with heterologous neutrophils for HBSS+ (white), mock-infected (blue bars) or RSV infected (red bars) cultures. Statistical comparison between all groups was performed using a paired t-test. Statistical significance is shown.

Neutrophils incubated with RSV infected ciliated epithelial cells released more active neutrophil elastase (NE) at 24h and 72h post-RSV infection (P<0.05) (**Figure 5C**). We found that the concentration of NE in apical surface media was significantly greater after 1h neutrophil exposure to RSV infected AECs at both 24h (P=0.01) and 72h (P=0.002) post infection with a mean±SEM of 54.5±13.5mU/ml, compared to 19.8±.4.4mU/ml in the mock-infected cultures at 24h post infection. To determine whether NE correlated with augmented azurophil granule release or was specific to NE alone, myeloperoxidase (MPO) activity was also assessed. Here we found significantly more MPO activity (P=0.003) after exposure of neutrophils for 4h to epithelial cells infected with RSV for 24h (P=0.006) or 72h (P<0.0001) (**Figure 5D**) with a mean±SEM of 1.0±0.0 OD at 24h, compared to 0.6±.0.1 OD in the mock-infected cultures. This was also significantly different after 1h at 72h (P<0.003) but not 24h (P=0.065) post-RSV infection.

RSV infection was also associated with greater release of active MMP-9 from neutrophils (**Figure 5E**) after 1h and 4h (P=0.001) in cultures infected with RSV for 24h; from 529.4±.21.9µg/ml in the mock to 949.3±.76.9µg/ml in RSV infected epithelial cells at 1h (P=0.02). There was no significance difference in MMP-9 concentrations between RSV and mock infected co-cultures at 72h post infection at either 1h (P=0.30) or 4h (P=0.07).

## DISCUSSION

We have shown that when RSV infected human primary nasal airway epithelial cells are exposed to neutrophils at physiological concentrations there is increased epithelial layer disruption, ciliary loss and less infectious virus in these cultures. This suggests that neutrophils are helping to eliminate viral infected cells and reduce viral spread. The airway epithelial damage that we observed, consistent with previous studies (20, 21), may be a necessary consequence of this anti-viral effect. We found that RSV infection without neutrophils did not reduce CBF, but did increase the number of cilia that presented with an abnormal beat pattern as early as 24h post-infection, which is similar to our previous findings (3, 5). Interestingly, the addition of neutrophils did not further increase ciliary dyskinesia.

The reduction in number of RSV infected epithelial cells following neutrophil exposure may result from neutrophil degranulation. Neutrophils are known to mediate direct antimicrobial effects and neutralize several influenza A, RSV and vaccinia virus strains through effector mechanisms including degranulation (22-24). We have shown that neutrophils exposed to RSV infected human primary airway epithelial cells have greater expression of the activation markers CD11B and MPO, and release greater amounts of NE, MPO and MMP9. It is recognized that the inflammatory processes in the airways of infants with RSV bronchiolitis are dominated by an intense neutrophil influx (7, 25) and that neutrophil products such as MPO and NE are released into the airway lumen (26). Indeed, the degree of neutrophilic inflammation correlates with disease severity in patients with RSV-induced bronchiolitis (27). Our study has shown that RSV infected epithelial cells increased NE and MPO and gelatinase (MMP-9) granule populations, at the same time as reducing neutrophil membrane integrity. This appears to be relevant to the pathophysiology of viral respiratory infections (28, 29). Likewise, both NE and MMP-9, which may be necessary for clearance of bacteria, are linked to airway damage and progression of cystic fibrosis (30). Our results are consistent with these clinical findings, suggesting that the cytotoxicity of neutrophil antimicrobial proteases may be important in viral clearance, but may also potentiate RSV-induced lung injury.

A surprising finding was that at 72h, but not at 24h, post RSV-infection, neutrophil exposure led to an increase in numbers of apoptotic neutrophils. This finding is consistent with a clinical study that found neutrophil apoptosis was accelerated in nasopharyngeal aspirates and peripheral blood of infants with RSV bronchiolitis (31). Neutrophil apoptosis is thought to be associated with loss of degranulation and other pro-inflammatory capacities (32). We found that RSV infected epithelial cells increased neutrophil degranulation in regards to increased neutrophil elastase (NE) and myeloperoxidase (MPO) activity in apical surface media at the same time as we detected the increased neutrophil apoptosis. These data suggest that within this model, there may be at least two subsets of neutrophils that respond differently to RSV infected airway epithelial cells. It is possible that this balance in neutrophil function could be a predictor of disease severity (See **Figure 6**).

**Figure 6.**
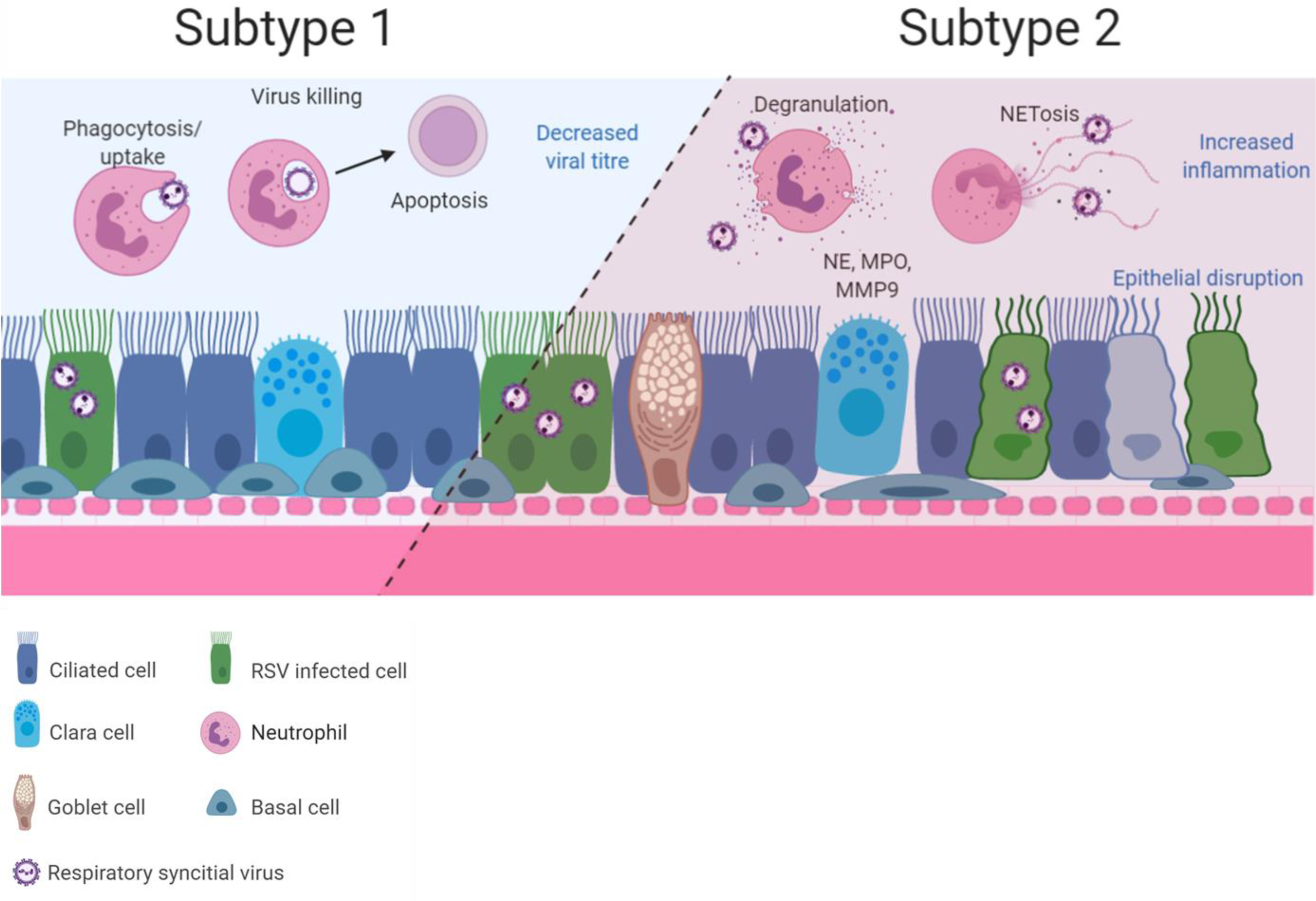
Diagram showing the two possible neutrophil subsets that maybe present during RSV infection of ciliated airway epithelial cells. This hypothesis may explain how simultaneous infection resolution and pathogenesis occurs in our model with results showing increased neutrophil activation, clearance of RSV and neutrophil apoptosis (Subtype 1). Our model also demonstrated neutrophil degranulation and release of proteases which are known to damage the epithelium (Subtype 2). Drawings created with BioRender.

One limitation to our study was the number of neutrophils used. The exact ratio of interacting neutrophils and airway epithelial cells in the lung of children with RSV bronchiolitis is unknown. We used the equivalent concentration of 5×10^6^/ml neutrophils, which is the upper limit of the number of neutrophils recovered from BAL of infants with RSV bronchiolitis (1.78±3.3×10^6^/ml) that was reported previously by McNamara and colleagues (7). This suggest that our data may indicate airway: immune cell interactions that occur in lungs of infants with large neutrophil infiltrate or severe RSV bronchiolitis. Another limitation of our study was that we exposed epithelial cells to naïve neutrophils directly isolated from peripheral blood. In the lungs, neutrophils are recruited to migrate from the basal sub-epithelial space across the vasculature and epithelium to the airways following RSV infection (33). In this context, the changes that we observed are all the more striking.

In conclusion, this study has revealed that neutrophils exposed to ciliated epithelial cell cultures infected with RSV have greater degranulation, deploying harmful proteins and proteases to the apical surface media and increasing the capacity for tissue injury. We have shown that neutrophils contribute to RSV-associated ciliary loss combined with epithelial damage, which is likely to result in reduce mucociliary clearance. These effects may contribute to viral clearance and provide important insights into the role of neutrophils in host response in the airway.

## ACKNOWLEDGMENTS

YD was a recipient of a Newton fellowship from The Academy of Medical Science (ref 0403). LR was a recipient of a Newton fellowship from The Academy of Medical Science (ref NIF004/1012) and Talent Training Programme of Children’s Hospital of Chongqing Medical University (class B abroad). CMS was a recipient of a grant from Wellcome Trust (212516/Z/18/Z). RLS was supported by the Great Ormond Street Children’s Charity (grant code W1802). This research was supported by the NIHR Great Ormond Street Hospital Biomedical Research Centre. Microscopy was performed at the Light Microscopy Core Facility, UCL GOS Institute of Child Health supported by the NIHR GOSH BRC award 17DD08.The views expressed are those of the author(s) and not necessarily those of the NHS, the NIHR or the Department of Health.

## AUTHOR CONTRIBUTIONS

All authors declare no conflicts of interest

YD: wrote funding application, conceived and designed the study, conducted the experiments, analysed data, and prepared the first draft of the manuscript.

JAH: assisted with the design of the study and data analysis.

ER: assisted with flow cytometry data analysis, and review of the manuscript.

LR: assisted with data analysis and review of the manuscript.

CMS: oversaw funding application, contributed to study conception, design, data analysis and interpretation, and final write-up of the manuscript.

RLS: oversaw funding application, contributed to study conception, design, data analysis and interpretation, and final write-up manuscript.

